# The hippocampus assumes a special role in supporting abstract concept representation

**DOI:** 10.1101/2023.07.04.547678

**Authors:** Alex Kafkas, Andrew R. Mayes, Daniela Montaldi

## Abstract

Unlike images, words are representational symbols. The associative details inherent within a word’s meaning and the visual imagery it generates, are inextricably connected to the way words are processed and represented. It is well recognised that the hippocampus associatively binds components of a memory to form a lasting representation, and here we show that the hippocampus is especially sensitive to abstract word processing. Using fMRI during recognition we found that the increased abstractness of words produced increased hippocampal activation and that critically this was independent of memory. Interestingly, word recollection produced hippocampal activation independent of word abstractness, while the parahippocampal cortex was sensitive to concrete word representation, independent of memory. We reason that the hippocampus has assumed a critical role in the representation of uncontextualized abstract word meaning, as its information-binding ability allows the retrieval of the semantic and visual associates that, when bound together, generate the abstract concept represented by word symbols. These novel findings not only offer insights for research drawing on word representation, memory, and hippocampal integrity, but critically, they throw important light on how the human brain may have adapted to encode and represent abstract words and concepts as they emerged in our language.

## INTRODUCTION

The perceptual features of objects around us, and scenes we are part of, can be encoded as visual representations using an approximation of a one-to-one mapping process. However, word-based information, which is a more recent addition to our information processing, cannot be meaningfully represented in this way. The perceptual features of a word (i.e., letters or sounds), unlike those of an object or scene, cannot generally form the basis of a representation that holds meaning or endures in memory. Instead the conceptual meaning, or semantics, as well as the perceptual details that a word brings to mind, are what form the basis of an enduring representation accessible to memory. Exactly how the brain might have adapted to support this remains unclear. Moreover, exactly how word information is represented in the brain can impact the type of memory generated and determine the specific brain regions, or networks, involved^1–4^. Indeed, words have been used in a vast number of studies exploring the neural basis of long-term memory and in most cases, findings from word-based studies have been generalised to all forms of information. Whether this generalisation is justified or whether words are processed differently remains to be established. Most central to this study is the consideration of the different evolutionary time over which language^5^ and perceptual processes emerged in the human brain, as we ask whether the brain’s memory systems have adapted to support memory for verbal information.

The overwhelming majority of evidence suggests that the hippocampus plays a selective and critical role in recall-based memory (or recollection)^6–10^ and in associating or binding together components into episodes^11–14^, while memory based on familiarity, absent of recall, is supported by the medial temporal lobe cortices^6,7,9,10,15^. Indeed, this evidence has led to the generally held view that these two kinds of memory are supported by contrasting computational algorithms and neural mechanisms^10,16^. However, one striking difference to this pattern is found when word stimuli are used (rather than objects, faces, or scenes). Then, more often than not, the hippocampus is reported to support familiarity as well as recollection^8,17–19^. We hypothesise that this difference, rather than presenting us with an unexplained conundrum, might be highlighting a unique role played by the hippocampus in the representation of words in memory^8^.

Important questions are, therefore, how might the hippocampus play such a special role in supporting memory for word-based information and what is it about words that makes them depend on the hippocampus. Answering these questions will help understand how, over time, the hippocampus has been adopted to support the representation of words, in a way that is not needed for other kinds of information. Using fMRI, we explored the neural mechanisms that support the representation and retrieval of word memory, while varying the abstractness of words. Abstractness describes the degree of visual imagery generated by a word and how tightly defined or not the meaning generated by a word may be^20,21^. We hypothesised that as memory for an abstract word cannot draw on a rich perceptual representation, and must require the bringing together, or association, of its key defining conceptual features, the neural mechanisms that support memory for words might differ as a function of word abstractness. We speculated that this adapted memory system might reflect the evolution of language in the brain from the more concrete (e.g., tools, animals etc) to the more abstract (e.g., conceptual, societal, emotional). To better understand the mechanism that might engage the hippocampus in word representation and memory, we exploited the algorithmic differences inherent in the familiarity/recollection distinction.

## RESULTS

Participants (N = 18; mean age = 22.78 years; SD = 2.21) encoded a series of abstract and concrete words (134 words for each type) outside the scanner, while making frequency judgements for each word (see Figure 1a and Methods). In the scanner, studied (268) and unstudied (134) words were presented and participants were trained to make recognition memory judgements for each word. An established memory protocol^6–8,22,23^ was used in which memory was examined by asking participants to discriminate recollected (R), familiar (on a rating scale from weak to strong familiarity) and new words (Fig. 1a). This allows the discrimination between those judgments that were based on memory for the stimulus word itself, with variable degrees of memory strength (from weak to strong), and those based on recollecting additional non-stimulus associative details from the encoding task^7,24^.

**Fig. 1.**
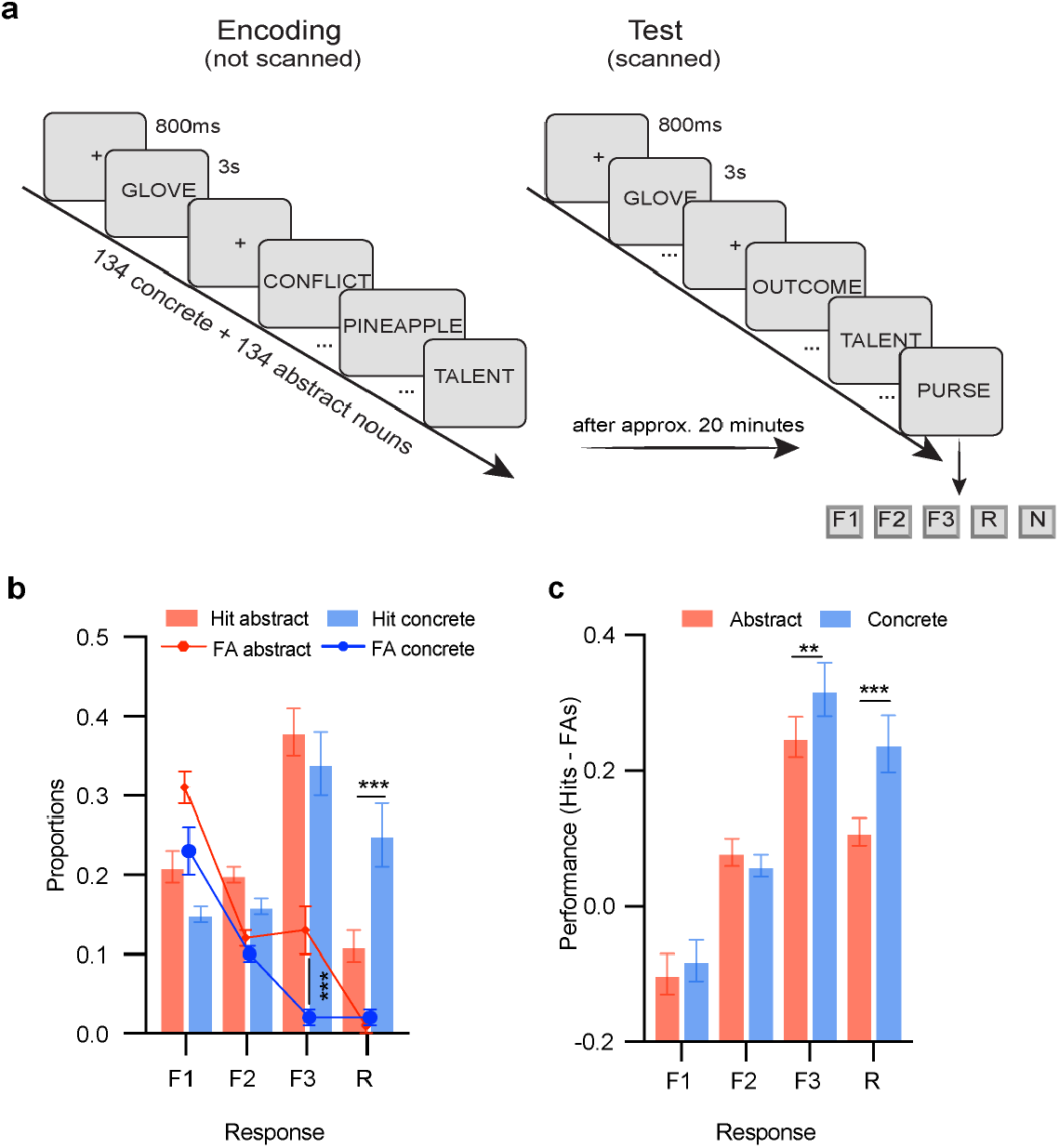
Design of the fMRI study and behavioural results. **a)** At encoding a series of abstract and concrete words were studied, while word frequency judgements were made. At test, while scanned, participants engaged in a recognition memory task in which previously studied words could be reported as recollected (i.e., retrieval of associated details related to the study episode), or familiar (i.e., words were recognised as being studied at encoding without triggering recall of additional details from study). Familiarity was reported using a rating scale (weak: F1, moderate: F2, strong: F3 familiarity), while separate and single responses were used to report recollected (R) and new/unstudied (N) stimuli. b) Proportions of hits (accurate old responses; bars) and false alarms (FA; incorrect old responses; lines) across the different memory outcomes plotted separately for abstract and concrete words. c) Memory performance for successful recognition of old stimuli (F1, F2, F3 and R) for abstract and concrete words, calculated as proportion of hits minus proportion of false alarms.

### Behavioural results

The proportion of trials for hits and false alarms (FA) across the different recognition responses are shown in Figure 1b, while memory performance is shown in Figure 1c, (response times and proportions across all memory types are presented in Supplementary Table 1). Abstract words, relative to concrete ones, were characterised by significantly increased FAs (*t*_17_ = 4.13, *p* = 0.001), increased F hits (*t*_17_ = 4.97, *p* < 0.001) and reduced recollections (*t*_17_ = 5.33, *p* < 0.001). Analysis of memory performance (hits minus FAs; Fig. 1c) showed advantage for concrete over abstract words (*F*_1,17_ = 9.01, *p* = 0.008, *η*^*2*^ = 0.35) owing to the increased FA rate to abstract words as described above (Fig. 1b). This was more pronounced in the case of strong familiarity and recollection (Fig. 1c; interaction *F*_2,34_ = 3.05, *p* = 0.06, *η*^*2*^ = 0.15; post-hoc abstract vs. concrete F1: *t* < 1; F2: *t*_17_ = 1.46, *p* = 0.16; F3: *t*_17_ = 3.01, *p* = 0.008; *t*_16_ = 4.55, *p* < 0.001).

### Neuroimaging results

#### Contrasting abstract-sensitive and concrete-sensitive brain regions

First, we explored the brain regions that increase their activity as a function of concreteness or abstractness, independent of participants’ memory judgment. A whole-brain analysis was performed on a trial-by-trial basis, treating each trial as a separate condition in the GLM, with the degree of abstractness as the parametric modulator (see Methods; findings in Fig. 2 and Supplementary Table 2). Activity in the bilateral hippocampus, the bilateral insula (BA13), the left middle occipital and lingual gyrus (BA19) and the middle cingulate (BA31) tracked increased word abstractness. On the other hand, decreases in abstractness (i.e., increases in concreteness) resulted in increased activity in the parahippocampal cortex (BA36), the precuneus (BA7/31), the superior medial and lateral prefrontal cortex (BA6/8) and the inferior parietal lobe (BA39/40).

**Fig. 2.**
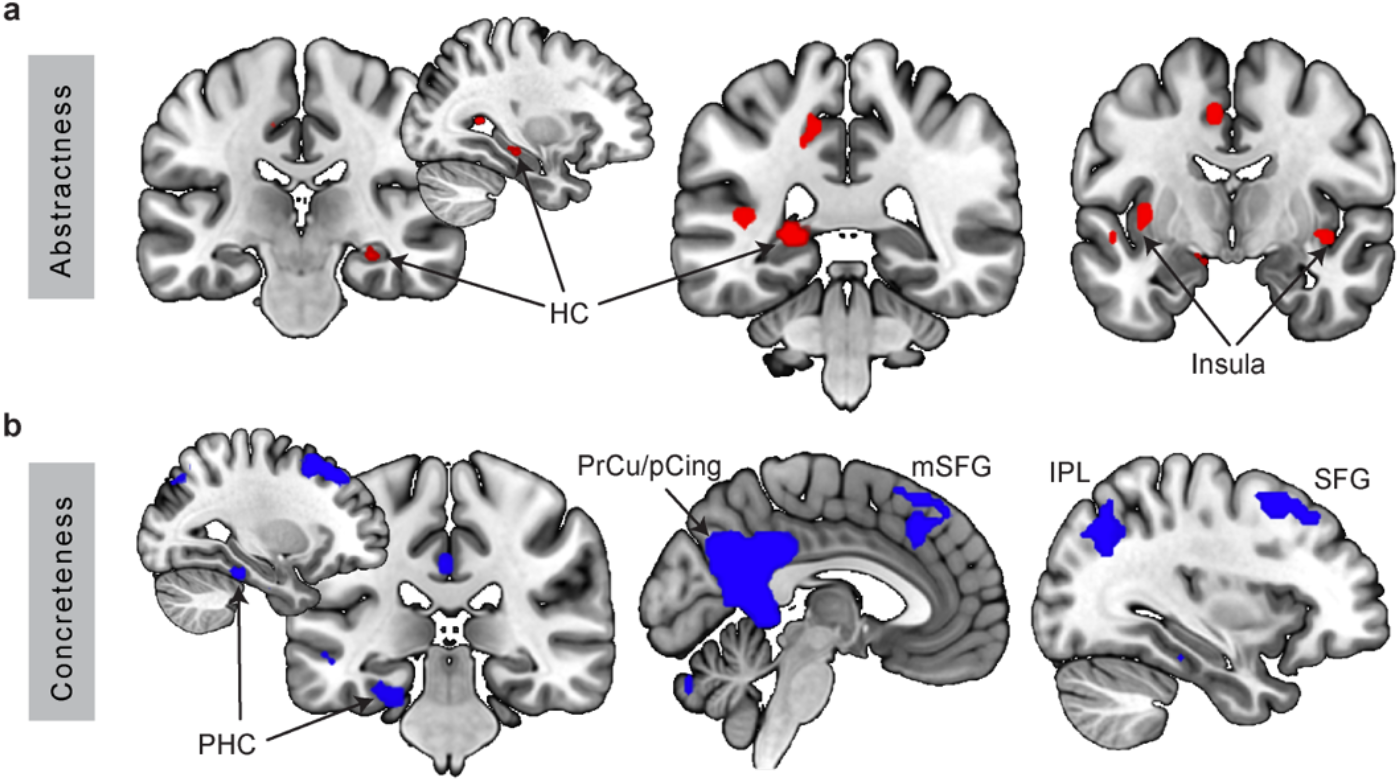
Whole-brain responses to degree of (a) abstractness (red) and (b) concreteness (blue) of the words. The illustrated regions are characterised by increased BOLD as a function of increased word abstractness (in red, upper panel) or increased word concreteness (in blue, lower panel) irrespective of kind of memory reported. HC = hippocampus; PHC = parahippocampal cortex; PrCu = precuneus; pCing = posterior cingulate; mSFG = medial superior frontal gyrus; IPL = Inferior parietal lobe; SFG = superior frontal gyrus. Activations are displayed at a voxel-wise *p* < 0.001 and are significant at a cluster-corrected FWE *p* < 0.05 determined via nonparametric permutations (all *t*s > 3.75).

#### Exploring abstract words and recognition memory in the hippocampus

As our data shows that the hippocampus is particularly sensitive to abstract words, we asked whether this sensitivity interacts with, or is driven by, or is indeed independent of the kind of memory experienced for the words. A series of analyses are described below that address this question separately for the two kinds of memory that support recognition. As recollection depends on, and reliably activates, the hippocampus^7^ if recollected abstract words should activate the hippocampus it would not be surprising. Therefore, we first sought to clarify the relationship between recollection and the hippocampal response to increased abstractness. We then explored this relationship with familiarity, a form of recognition not generally believed to engage the hippocampus, except, perhaps, when word stimuli are used, as alluded to in the Introduction.

#### The hippocampus is sensitive to word recollection

The analyses so far have established that the hippocampus responds to word abstractness irrespective of memory outcome. But does this hippocampal effect interact with ongoing recollection, which involves the recall of non-stimulus details from encoding? Whole brain responses to recollection relative to misses and strong familiarity (F3) for both abstract and concrete words are shown in Figure 3a-b (see also Supplementary Tables 3 and 4). A univariate analysis indicated that recollection relative both to misses and to F3 produced activations within the hippocampus independent of abstractness (Fig. 3a and 3b). A separate parametric analysis, with abstractness as a parametric modulator on recollected trials only, did not produce any hippocampal effects (or effects in any other parts of the brain) either for increased abstractness or increased concreteness. Finally, MVPA classification analysis showed that activation within the hippocampus (bilaterally and separately for left and right) significantly discriminated recollected events from missed and F3 trials (Fig. 3c and 3d); see also data in Supplementary Table 5). These findings clearly reveal reliable hippocampal activations for recollection, which while diagnostic of recollection, are independent of abstractness.

**Fig. 3.**
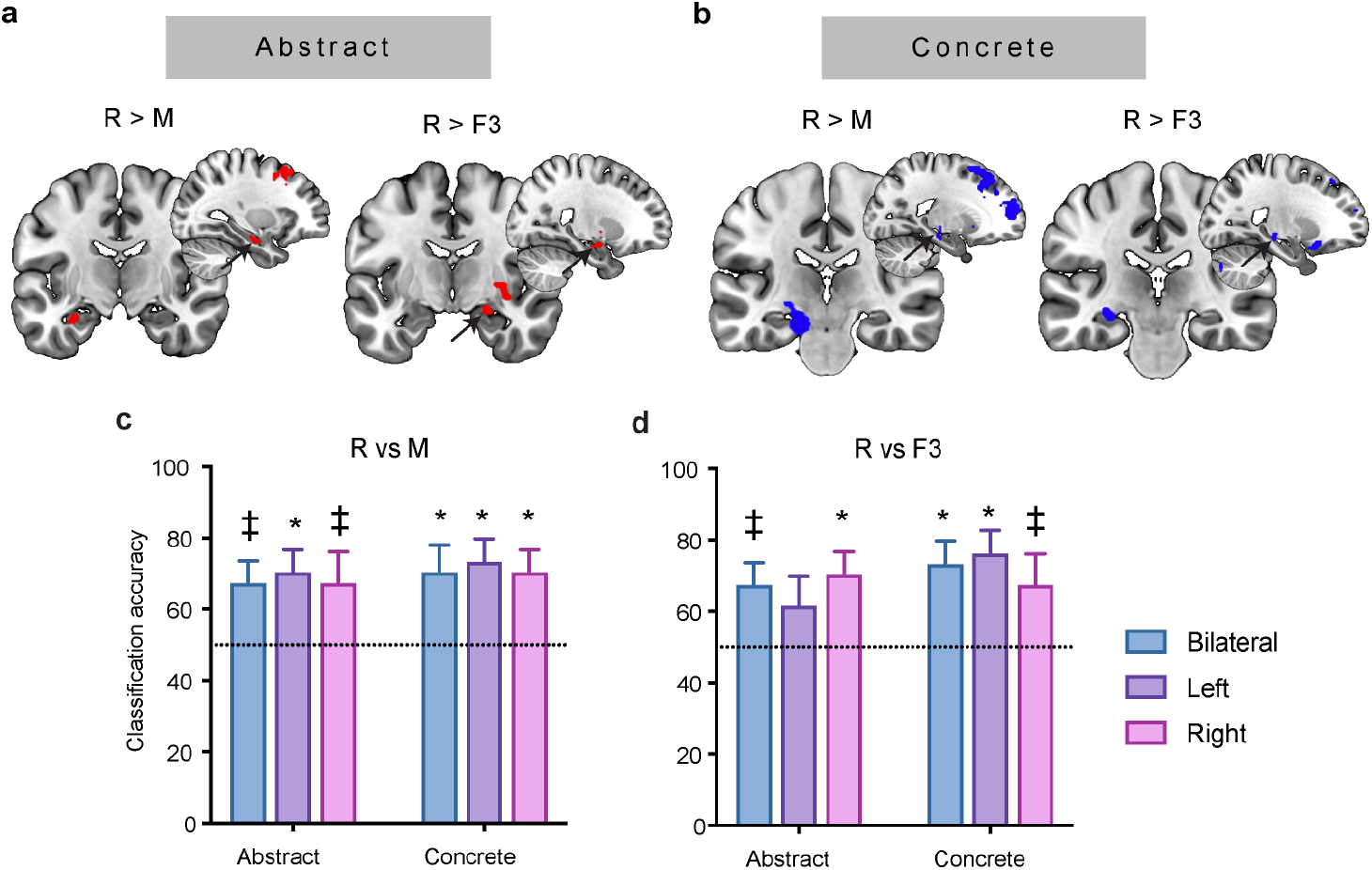
Hippocampal responses to recollection of abstract and concrete words: Activations are strongly diagnostic of memory. **a-b)** Hippocampal activations for abstract (red; a) and concrete (blue; b) words reported as recollected relative to misses and F3 responses. R > M for abstract: MNI: -30 -13 -20; R > M for concrete: MNI: -15 -25 -17, cluster includes hippocampus and parahippocampal cortex. R > F3 for abstract MNI: 21 -7 -14, (cluster centred within the amygdala including 9 voxels in the anterior hippocampus); R > F3 for concrete MNI: -24 -28 -5). Activations are displayed at a voxel-wise *p* < 0.001 level and are significant with a cluster-corrected family-wise error (FWE) *p* < 0.05 determined via nonparametric permutations. **c-d)** Classification (MVPA) outcomes within the hippocampus showing that hippocampal activity discriminates recollections (relative to misses and F3) for both abstract and concrete words. In separate analyses, binary classification success was calculated for recollected versus missed (c) and recollected versus F3 trials (d). Significance of classification success from activation data was assessed based on permutation testing with 5000 permutations. * *p* < 0.05; ‡ *p* = 0.056/0.054 (trend). Data (percent accuracy and *p*-values) are also presented in Supplementary Table 5. Error bars indicate the standard error of the mean across participants.

#### The hippocampal response to word familiarity is driven by abstractness

We have established that the hippocampus responds to word abstractness irrespective of memory outcome, and that the hippocampal role in recollection is reflected in its activation for word recollection, independent of abstractness. Next, we turned to explore hippocampal sensitivity to abstract and concrete words when memory was supported by familiarity. First, parametric analyses explored how rated familiarity strength modulated regional brain activity, separately for abstract and concrete words (see Methods). Hippocampal and caudate nucleus activity were uniquely modulated by abstract word familiarity, while parahippocampal cortex (BA 35) activity was uniquely modulated by concrete word familiarity (Suppl. Table 6). Exclusive masking methodology (see Methods) confirmed that these parametric familiarity responses were selective for abstract words (caudate nucleus and hippocampus) and concrete words (parahippocampal cortex; Fig. 4a, 4b and Suppl. Fig. 1). In addition to these abstract-unique and concrete-unique activations, these analyses also revealed a network of somewhat overlapping extra-MTL brain regions sensitive to abstract and concrete word familiarity strength (see Suppl. Table 6 and Suppl. Fig. 2).

**Fig. 4.**
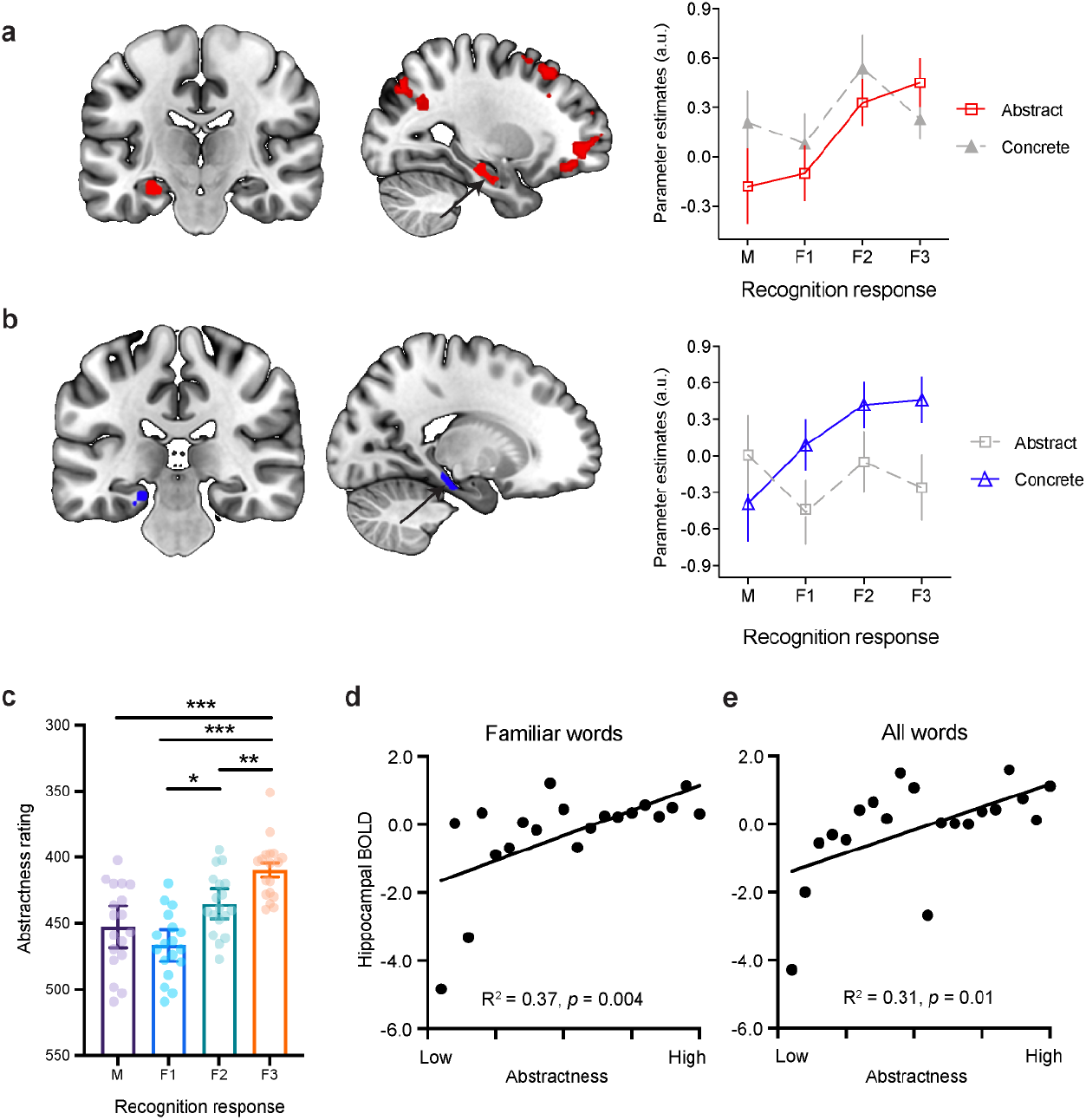
Activation increases in the MTL as a function of familiarity memory. **a)** Hippocampal activity (MNI: -30 -22 -14) is modulated by abstract word familiarity strength (surviving exclusive masking by concrete word familiarity). **b)** Familiarity response in the left parahippocampal cortex (MNI: -22 -27 -20) selective for concrete words (surviving exclusive masking by abstract word familiarity). Activations are displayed at a voxel-wise *p* < 0.001 and are significant at a cluster-corrected family-wise error (FWE) *p* < 0.05 determined via nonparametric permutations. The parameter estimates plotted in red (a) or blue (b) indicate which familiarity parametric response was selectively significant in each region – either familiarity response to abstract or concrete words; the non-significant parametric response is plotted in gray. **c)** Mean word abstractness rating for each recognition response category (smaller numbers on y-axis indicate greater abstractness) demonstrate that increased abstractness is associated with increased familiarity strength. **d) and e)** Regression analysis between hippocampal activity in the cluster presented in (a) and the degree of abstractness (from highly concrete to highly abstract words) for words judged as familiar (d) and all words (e) irrespective of memory outcome. Note: M=misses, F1 = weak, F2 = moderate, F3 = strong familiarity hits. \*\*\**p* < 0.001; ** *p* < 0.01; * *p* < 0.05. All error bars show the standard error of the mean.

#### Abstractness drives the hippocampal activity independent of familiarity

A regression analysis of the familiar words showed that the degree of hippocampal activation (from a 4mm sphere centred around -30 -22 -14 as in Fig. 4a) significantly tracked the degree of word abstractness with lower activation for the most concrete words and higher activation for the most abstract ones (R^2^ = 0.37, *p* = 0.004; Fig. 4d). This was also true when abstract (R^2^ = 0.57, *p* = 0.011) and concrete (R^2^ = 0.48, *p* = 0.025) words were assessed separately. Furthermore, BOLD activity in the hippocampus tracked the degree of abstractness across all words encountered at test, independent of what response was given and of whether previously encoded (Fig. 4e), with greater activation generated by the most abstract words (R^2^ = 0.31, *p* = 0.01). Again, this was also true when abstract and concrete words were examined separately (concrete: R^2^ = 0.76, *p =* 0.001; Abstract: R^2^ = 0.49, *p =* 0.02). Therefore, the hippocampus responds reliably to the degree of abstractness of words, and this is true not only in the case of words reported as familiar but for any word, irrespective of memory status or memory response.

To identify any relationship between abstractness and familiarity ratings, the average abstractness rating of words rated as F1, F2 and F3 (as well as misses) was calculated for each participant (Fig. 4c). The mean abstract rating increased with increased familiarity memory strength (*F*_3,51_ = 17.28, *p* < 0.001, *η*^*2*^ = 0.50) with mean abstractness rating being significantly higher for F3 compared to each of the other levels (all *p*s < 0.01; see Fig. 4c). This strongly suggests that the familiarity-related activation in the hippocampus (Fig. 4c) was driven predominantly by abstractness.

Furthermore, with MVPA we examined whether cluster activity within the anatomical region of the hippocampus (bilateral and left/right separately) could predict the accuracy with which familiar words were discriminated from missed words. As shown in Figure 5a, activity within the bilateral hippocampus did not discriminate familiar from missed abstract words (accuracy = 38.9%, *p* = 0.84) or concrete words (accuracy = 50%, *p =* 0.57), and similar results were observed when examining left and right hippocampus separately (Fig. 5). However, when examining the hippocampal activity classification performance for all abstract and concrete words (irrespective of memory status or response), classification accuracy was above chance for abstract (accuracy = 77.8%, *p* = 0.014) but not concrete words (accuracy = 50%, *p =* 0.53; Fig. 5b). The effect for the abstract words was evident in the left hippocampus (accuracy = 72.22%, *p* = 0.03) but not the right hippocampus (accuracy = 55.6%, *p* = 0.41). Therefore, this MVPA analysis demonstrates that hippocampal activity does not reflect familiarity memory but instead, is sensitive to the abstractness of the words.

**Fig. 5.**
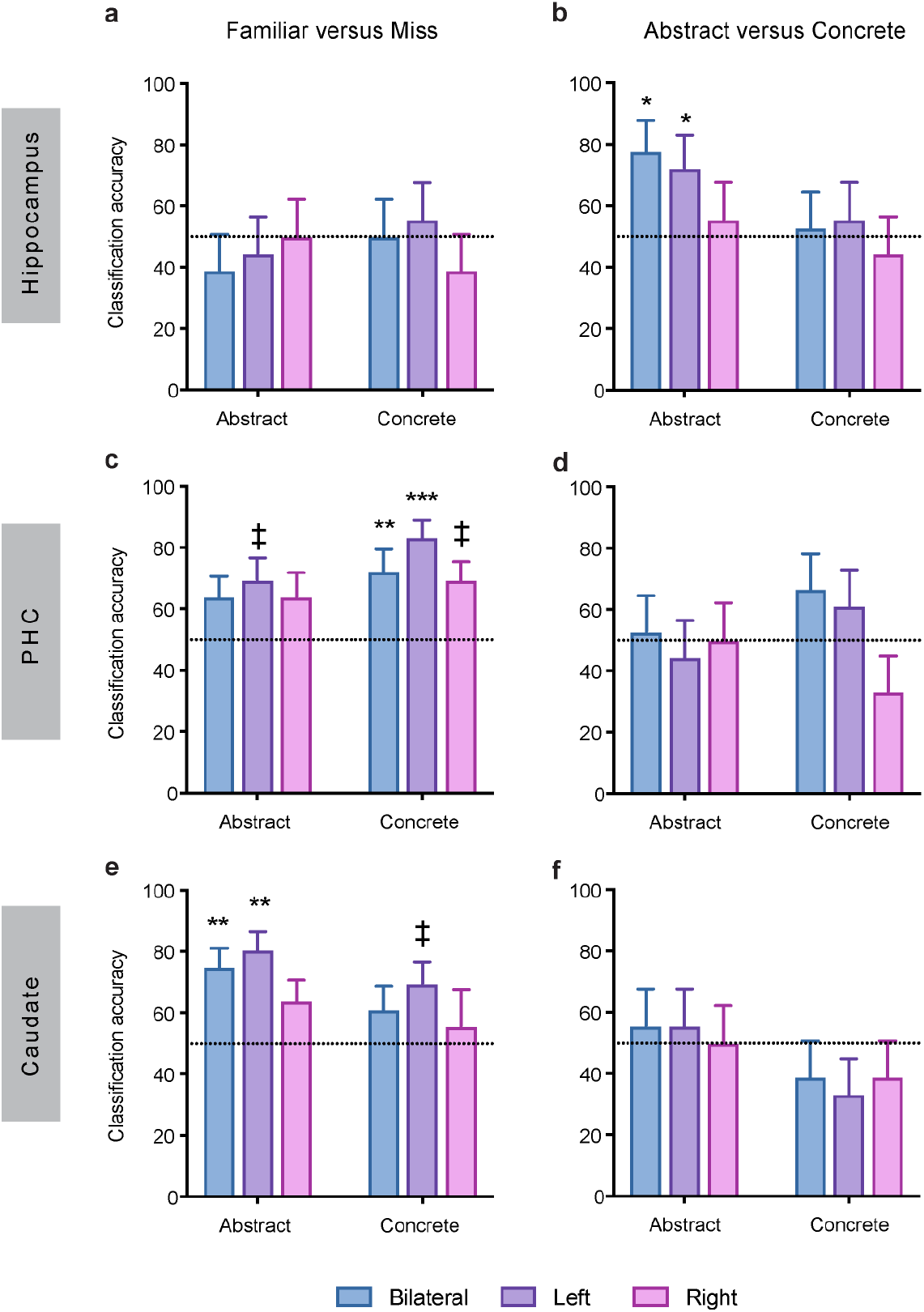
**Classification (MVPA) outcomes within the hippocampus (a, b), the parahippocampal cortex (PHC; c, d) and the caudate nucleus (e, f) for abstract and concrete words judged as familiar (a, c, e) and all word stimuli irrespective of memory status or response (b, d, f)**. For the familiar word analyses, binary classification success was calculated compared to misses (i.e., familiar abstract words vs. missed abstract words and familiar concrete words vs. missed concrete words). For the abstract and concrete analyses, binary classification success was calculated versus baseline activity (i.e., abstract words vs. baseline and concrete words vs. baseline). Significance was assessed based on permutation testing with 5000 permutations. * *p* < 0.05; ** *p* < 0.01; *** *p* < 0.001; ‡ *p* = 0.056/0.054 (trend). Error bars indicate the standard error of the mean across participants.

Further support for the selectivity of this hippocampal sensitivity to word abstractness was provided by an MVPA analysis of the caudate nucleus (Fig. 5e-f) and the parahippocampal cortex (Fig. 5c-d), regions that showed selective univariate responses to familiar abstract (caudate) and familiar concrete (parahippocampal cortex) words (Fig. 4b and Suppl. Fig. 1). These analyses showed (see Fig. 5 and Supplementary Analyses) that unlike the hippocampus, activity within these regions was driven predominantly by familiarity memory and not by the degree of abstractness of the words.

## DISCUSSION

Words are symbols used to express concepts from the most concrete to the most abstract. Their functional emergence and subsequent evolution must have triggered a degree of brain adaptation reflecting the evolution of human thought and language, which likely originated with the most concrete concepts, such as those describing people, objects, and animals. Only much later will human thought have evolved to require expression of highly abstract concepts. Thus, it would not be surprising if the brain had to adapt its word representational system, to readily support abstract concepts. We believe the current study provides exciting new evidence in support of this view. Specifically, this study was motivated by the observation that the hippocampus appears sensitive to word familiarity but not familiarity for other types of information^7,8^. We asked what might be particular about words that they draw on the hippocampus to support a kind of memory (i.e., familiarity) which is described as being largely non-associative, and almost universally, as non-hippocampal (for discussion see^9,25^). We therefore explored whether the nature of word representation sometimes requires a specialised system whereby the representations of words, or some features of words, are supported by a hippocampal mechanism.

While as expected, we found the hippocampus to be highly sensitive to word recollection (whether the word was high or low in abstractness), we strikingly also found that hippocampal activity systematically and reliably tracked the degree of abstractness of a word independent of memory. Furthermore, our data shows that abstractness, rather than familiarity, modulates, and is predicted by, hippocampal activity. This suggests that, at least under some conditions, when words are encountered, the hippocampus is recruited to help form the associative representation of words that are somewhat, or very, abstract, irrespective of whether that representation is experienced as a memory or not. We believe that this particular hippocampal role aids the creation and evaluation of a representation by retrieving the semantic and conceptual associates of a word without triggering recollection of the word itself. Indeed, it has been shown that hippocampal damage may result in impaired access to semantic knowledge related to vocabulary depth and richness^26, see also 27^.

This proposed hippocampal function is consistent with the established role of the hippocampus in forming and retrieving associative memories^12–14,28^. The same associative mechanisms are, therefore, likely to facilitate the processing of verbal information when abstract word content is encountered. Indeed, according to the semantic memory framework, proposed by Crutch and Warrington^29^, abstract words are organised in terms of associative properties and relationships, and more so than concrete ones. They argue that the representational systems that support abstract and concrete word meanings, have qualitatively different properties. Consistent with this is the argument that abstract words are relatively unconstrained perceptually or spatially, and are linked with diverse contexts relative to concrete words^20^. Indeed, from a linguistic perspective, it is argued that abstract words show high levels of semantic variability and complexity and are associated with a wide variety of linguistic and semantic contexts^30,31^, including for example, those characterised by emotionality, social interaction, morality, time, and space^21^. From a neural perspective, therefore, since the substrates of abstract concepts are somewhat determined by their semantic content, their neural bases might be expected to vary more than for concrete words^32^. Indeed, brain activation prediction models have been found to successfully predict concrete words, but fail to predict abstract words^33^, which is consistent with the limited agreement between participants regarding the definitions of abstract, compared to concrete, words^34^.

The difference between our experience of concrete and abstract words is also reflected in the behavioural performance of our participants and explains why, what might at first appear to be a hippocampal sensitivity to familiarity, is instead driven by its sensitivity to abstractness. Critically, the increase in strong familiarity, and false alarms, reported by participants in response to abstract compared to concrete words, indicates that participants were more likely to attribute feelings of memory to familiarity than recollection in the case of abstract words, even when memory was illusory. Furthermore, the strength of reported familiarity increased with increasing abstractness (Fig. 4c), while hippocampal activity did not discriminate familiar from unfamiliar (i.e., forgotten) words, but abstract from concrete ones (Fig. 5a-b). Here we propose that the abstractness of a word triggers the spontaneous generation of hippocampally-dependent associations, that when bound together produces the word’s conceptual representation. We suggest that this process draws on the same hippocampal mechanism that supports episodic associative memory^8,12^, and it draws on it increasingly, with increasing levels of abstractness. We therefore argue that since the level of activity in the hippocampus reflects the amount of information it is associating^8^, the more abstract a word, the more its representation depends on the hippocampus. In turn, as both hippocampal activity and associative processing increase, so does the richness and familiarity of the representation experienced. Furthermore, the product of the hippocampal signal, while actually reflective of associative processes, is interpreted by participants as familiarity. This signal generates the concept that the word itself represents, and not extra-stimulus associative detail, memory for which would be supported by recollection.

In contrast, the processing of concrete words relies more directly on visual information^35^ involving the retrieval of stored representations of the visual and conceptual features that concrete words represent^35–37^. As the parahippocampal cortex has a well-recognised role in processing complex visual information (e.g.,^38,39^), its activation for concrete words found in the current study is not surprising. Indeed, the parahippocampal activity, tracks the concreteness of words, independent of memory. Importantly, it also predicts familiarity, which likely reflects the contribution detailed visual information makes to the richness of a representation and the subsequent feeling of familiarity attributed to it. Therefore, we argue that these two MTL structures fulfil very different word-related functions; the hippocampus supports abstract word representation, as well as word recollection, while the parahippocampus supports concrete word representation, as well as word familiarity, but not word recollection or abstract word representation. Therefore, the functions of these structures combine to produce rich word representations but diverge to support different components of word recognition memory.

While we interpret the hippocampal sensitivity to word abstractness as independent of memory, the experiment itself involved a memory paradigm, and therefore participants were engaging in memory judgements, irrespective of whether words were identified as old or new, correctly, or incorrectly. The important implication here is that during scanning, all words were subjected to a memory evaluation which will have triggered mechanisms that either assessed the familiarity of each word or retrieved word representations from memory. Critically, however, even words that were not in memory, (i.e., not studied prior to the scan), showed hippocampal sensitivity to abstractness. So, while this effect reflects not only the formation of word representations, it also reflects the detailed evaluation of a word representation. This is important to highlight because it might be that under some conditions, the use of abstract words can be achieved with limited, or no hippocampal engagement, but perhaps when the generation of the representation in its richest form is required (perhaps to maximise memory accuracy), then the hippocampus might become critical. These findings have important implications for our understanding of memory and amnesia and potentially also our ability to detect early hippocampal deterioration (as seen in some dementias). Therefore, future research may explore differences in processing abstract (compared to concrete) in patients with hippocampal lesions, mild cognitive impairment, and healthy ageing.

Our novel findings reflect an adaptation of a memory system originally optimised for more concrete representations, where an approximate one-to-one mapping between object and representation can be supported by non-hippocampal structures in the ventral visual stream. We suggest that as human language has evolved from that dominated by the more basic object representations (tools, objects, faces), to incorporate more abstract concepts, generally associated with social organisation, morality, and emotion, our brains have adapted to support their representation. This proposed adaptation draws on the hippocampus and its highly specialised associative mechanisms. Without these mechanisms we predict that the brain is only able to robustly represent abstract concepts and words in their simplest, or most degraded form.

Taken together these data and their interpretation contribute important new ideas about how words and concepts are represented in the brain, and consequently how the neural bases of memory for words might be distinct from that of memory for other forms of information (e.g., objects, scenes, faces). Finally, this research offers highly novel, but compelling evidence that our representational system has likely adapted to address the needs of a more sophisticated, complex, and abstract language.

## METHODS

### Participants

In total, 21 participants gave informed consent and participated in the study. From these, 3 participants were excluded from the analyses; 1 due to a technical problem preventing the recording of fMRI data at retrieval and another 2 due to chance performance in the memory task. Therefore, the final sample included 18 participants (4 male), with a mean age of 22.78 years (SD = 2.21). The sample size was informed by our previous experiment utilising word stimuli with a similar experimental procedure^8^. Also, a power calculation (using NeuroPower; URL: http://neuropowertools.org/) using data from our previous study, determined that for 18 participants we gain a power of 60%, which is considered moderate and therefore acceptable for the aims of the present study. All participants were native English speakers, right-handed, neurologically healthy and had normal (uncorrected) vision. The study received ethical approval from The National Research Ethics Service (North West-GM South) and each participant was paid £20 after completing the testing session.

### Materials

Stimuli consisted of 402 word nouns (plus 14 words for practice) obtained from the MRC Psycholinguistic Database^40^. Abstract words were selected to have a concreteness rating lower than 400 (range = 204–371; mean = 309, SD = 37), while concrete words were selected to have a concreteness rating higher than 500 (range = 516–653; mean = 575, SD = 29). Overall concreteness rating was significantly different between abstract and concrete words (*t*_400_ = 79.71, *p* < 0.001). Concrete and abstract words were selected to have MRC familiarity ratings of at least 400 (abstract range: 420 – 612; concrete range: 421 – 657) and the two word groups were matched for familiarity rating (*t*_400_ = 1.11, *p* = 0.27), Kucera-Francis written frequency scores (*t*_400_ < 1) and word length (range = 5 – 9 letters).

### Experimental Procedure

Each experimental session consisted of two phases: an encoding phase completed outside the MRI scanner and a retrieval phase completed in the scanner (Figure 1a). At encoding participants studied a series of 268 words, randomly selected from a pool of 134 concrete, and 134 abstract. Each word was presented centrally, in black letters (font: Arial 25pt) on a white background for 3s. For each word participants were asked to make a frequency decision indicating how often they encounter each word in their everyday life (either spoken or written) using a binary decision (“quite often” or “not so often”). The selection of the encoding task was informed by previous unpublished piloting, which established that this task results in a good response spread across the familiarity memory rating scale and overall adequate memory performance. Three other encoding tasks (involving pleasantness, alphabetic and semantic decisions) were rejected as resulted in either poor overall memory performance (alphabetic decision) or in memory decisions restricted to the high end of the rating scale (either strong familiarity or recollection; pleasantness and semantic decision tasks).

After the encoding block and before starting the MRI scan, participants were trained in using a modified version of the remember/know procedure^6,7,12,22,24^. This entailed discriminating instances of familiarity, using a 3-point rating scale (from weak to strong familiarity: F1 = weak; F2 = moderate; F3 = strong familiarity), instances of recollection (R) and new stimuli (N). Participants were instructed to provide a familiarity response when they felt that they had encountered a stimulus at study, but report words as recollected if they spontaneously recalled additional associative information from the study episode in relation to a stimulus. After the training and to ensure understanding of these instructions, participants were asked to explain familiarity and recollection, to provide examples from their own experience and to ask questions about these two kinds of memory. A practice block that resembled the retrieval task was completed outside the scanner and one more practice was given in the scanner before the main task.

At retrieval, inside the scanner, participants were presented with 268 abstract and concrete words from encoding along with 134 new foils (67 concrete and 67 abstract). The fMRI data were collected and analysed from this phase and the session was divided into 2 functional runs. Each word was presented for 3s and participants were asked to use a 5-button MR-compatible button box to decide whether each stimulus is familiar, using three levels of increasing familiarity (i.e., F1, F2 or F3), whether it is recollected or new. Three fingers on one hand were used to select the familiarity responses and two fingers on the other hand were used for the extra two options (R and N). This allocation to the right or left hand were counterbalanced across participants.

Each word trial was followed by a fixation cross presented for 800ms and the selection of the type of the presented word in each trial (abstract or concrete) was random. Intermixed with the main events a set of 81 implicit baseline (null) events were also presented to ensure effective jittering and to provide baseline BOLD measures. During the fMRI session, participants were provided with earplugs and ear protection headphones and soft pads were used to stabilise their head to prevent movement artefacts.

### fMRI data acquisition and pre-processing

A 3T Philips Achieva scanner was used to acquire the MRI data. Functional data (gradient echo-planar images; EPI) were acquired using the blood oxygenation level dependent (BOLD) contrast and a total of 735 volumes (TR = 2.5s; TE = 35ms; 40 slices per volume; matrix size = 96 x 96; voxel size = 2.5 x 2.5 x 3.5 mm) were recorded for each participant across three sessions covering the whole brain. T1 high resolution images were also collected from each participant at the beginning of each session, before the functional blocks (180 slices; voxels size = 1mm isotropic; matrix size = 256 x 256).

The ArtRepair software (http://cibsr.stanford.edu/tools/human-brain-project/artrepair-software.html) was used to examine data quality of the EPI time-series and residual movement parameters were used in the GLM models (see below). SPM8 (Statistical Parametric Mapping, Wellcome Trust Centre for Neuroimaging; http://www.fil.ion.ucl.ac.uk/spm/) was used for data pre-processing and analysis. Data pre-processing included realignment of the EPI data to the mean image using a standard six-parameter rigid body transformation, reslicing using sinc interpolation and slice timing correction (to the middle slice). Each subject’s high resolution T1 image was coregistered to the corresponding mean EPI image. Finally, the coregisterd T1 and functional images were spatially normalised to MNI space using the DARTEL tool in SPM8^41^. DARTEL normalisation has been shown to provide improved realignment of the MTL structures across multiple participants when a whole-brain analysis is conducted^42^. After spatial normalisation the functional data were resliced to 3mm isotropic and were spatially smoothed using a 5mm isotropic full width half maximum (FWHM) Gaussian kernel.

### Univariate fMRI analyses

The pre-processed fMRI data were analysed using the general linear model (GLM) separately for each individual at the first level analysis. The onset (and the duration) of each event of interest was convolved with the canonical haemodynamic response function. For each participant, three models were defined, one trial-specific, one memory categorical and one memory parametric (see below). Regressors of no interest were also modelled and included trials with no behavioural response, the six movement parameters after realignment for each functional run and residual movement artefacts identified from the ArtRepair tool. A high-pass filter of 128s was applied to the data to remove low-frequency noise.

To explore BOLD responses as a function of stimulus abstractness irrespective of the type of memory response accompanying each word, a parametric model was constructed for each participant modelling all retrieval trials as separate conditions (trial-specific model). In this model, the abstractness rating of each word (obtained from the MRC Psycholinguistic database) was used as a covariate and was convolved with the trial-specific HRF. First-level parametric *t* contrasts were created for linear increases as a function of increased concreteness or linear decreases as a function of increased concreteness mapping therefore brain responses to increased abstractness. Each of these first-level t-contrasts were used in the second-level (group) analysis implemented as one-sample t-tests.

For exploring the memory interaction with abstractness, in the categorical model separate regressors for each condition of interest were defined for each participant. These consisted of all the potential memory outcomes separately for the two types of words (F1_abstract, F1_concrete, F2, abstract, F2_concrete, F3_abstract, F3_concrete, R_abstract, R_concrete, CR_abstract, CR_concrete, M_abstract, M_concrete, FA_collapsed). Trials with no responses were also modelled as conditions of no interest. In order to explore modulation of brain activity by familiarity strength for abstract and concrete words a parametric model was constructed for each participant^43^. In this model, familiarity hits (i.e., old stimuli reported as familiar) were specified as separate conditions, and the reported strength associated with each familiarity response was used as a covariate and was convolved with the onset-specific HRF. Two parametric conditions were specified like this for the two word types (abstract and concrete words). In each parametric condition, four levels of strength were specified with misses (old stimuli deemed new) used as the level with zero familiarity (*F*_0_), while F1, F2 and F3 responses were used as reflecting increasing levels of familiarity (from weak to strong). At the first (subject) level parametric *t* contrasts were created for linear (monotonic) increases or decreases in activity across familiarity strength for abstract and concrete words (separately) and each of them were used in the second level (random effects) analysis implemented as one-sample *t*-test. Non-linear (quadratic) effects were also modelled to capture residual non-linear effects, but as these did not produce additional activations (not already included in the linear contrasts), they are not reported separately.

Regions of common modulation by familiarity strength for abstract and concrete words were examined using conjunction analysis based on the conjunction null hypothesis^44^. The parametric contrasts were also analysed using exclusive masking to explore activations unique to each type of stimulus (i.e., parametric familiarity for abstract by parametric familiarity for concrete words and vice versa; exclusive mask threshold: *p* < 0.05). This analysis reveals the brain regions that are modulated by familiarity strength for one type of words but not for the other (i.e., uniquely characterise familiarity for either abstract or concrete words). Recollection-related activations were also explored in the whole brain by contrasting recollections to abstract and concrete words with misses and F3 responses (i.e., recollections to abstract words: R_abstract_ > M_abstract_ and R_abstract_ > F3_abstract_; recollections to concrete words: R_concrete_ > M_concrete_ and R_concrete_ > F3_concrete_). Significance in all analyses was determined at a cluster-corrected family-wise error (FWE) *p* < 0.05 determined via nonparametric permutations as implemented within the Statistical NonParametric Mapping toolbox (SnPM 13; URL: http://warwick.ac.uk/snpm).

### Regression analysis

Using the trial-specific GLM model, activation data (parameter estimates) for each trial and each participant were extracted from a hippocampal cluster identified in the whole-brain parametric analysis using a mask of 6mm sphere around the MNI peak (−30 -22 -14). Linear regression analyses were performed between the hippocampal activation data and degree of concreteness/abstractness (MRC Psycholinguistic rate of concreteness) of each word for the familiar trials (i.e., words which participants reported as familiar) as well as for all the words irrespective of participants’ memory outcome.

### Multivoxel pattern analysis

Multivoxel patterns analysis (MVPA) was also conducted to further characterise the extent to which hippocampal activity successfully predicted (or classified) abstract and concrete words based on their reported familiarity or their property as abstract or concrete. Similar analyses were also conducted for the parahippocampal cortex and caudate nucleus, as these structures were also found to respond to stimulus reported familiarity (the latter are presented in Suppl. Analyses). Anatomical masks of the bilateral hippocampus, the bilateral parahippocampal cortex and the bilateral caudate nucleus were used as Regions of Interest (ROIs) derived using the PickAtlas Toolbox^45^. The Pattern Recognition for Neuroimaging Toolbox (PRONTO, http://www.mlnl.cs.ucl.ac.uk/pronto/^46^) was used for the MVPA analysis. Classification accuracy in the ROIs was explored for familiar (collapsed F1, F2, F3 responses) abstract and concrete words relative to misses for each word type (i.e., familiar abstract vs. missed abstract and familiar concrete vs. missed concrete) using a set of binary support vector machine (SVM) classification procedures. Furthermore, classification success within the ROIs was also assessed for abstract vs. concrete words irrespective of the behavioural response associated with each response using a binary SVM algorithm (i.e., abstract words vs baseline and concrete words versus baseline). The classification analyses were performed within the bilateral masks and separately for left and right lateralised masks. In the analyses the data were mean centred and a leave-one-subject-out cross validation method was adopted to perform group analyses. Statistical significance of the classification outcomes (accuracies) for each model within the ROIs was tested using non-parametric permutations with 5000 iterations.

## Data availability

Study material are available from the corresponding author on reasonable request.

## Acknowledgements

This work was supported by the Wellcome Trust (grant number: 094597/Z/10/Z).

## Author contributions

The experiment was conceptualised and designed by all authors. A.K. collected, analysed and visualised the data. Data interpretation and manuscript writing was carried out jointly by all authors.

