## Supplementary Material for "The hippocampus assumes a special role in supporting abstract concept representation"

### Supplementary Analyses/Results

#### Multivoxel pattern analysis in the caudate nucleus and the parahippocampal cortex

In order to further explore the selectivity of the MVPA effect within the hippocampus (as reported in the main text and shown in Fig 5), MVPA methodology was also adopted to explore classification outcomes within the caudate nucleus and the parahippocampal cortex. These two regions showed selective univariate parametric response to familiar abstract words (caudate; Supplementary Fig. 1) and familiar concrete words (parahippocampal cortex; Fig. 4b), therefore it is relevant to explore whether these responses truly reflect sensitivity to familiarity memory or they are confounded by the abstractness of the words as was the case for the hippocampus. These findings are also important for the selectivity of the identified effects in the hippocampus in response to familiarity. In other words, is familiarity-related univariate activation truly diagnostic of familiarity in other brain regions?

As shown in Figure 5e, activity within the bilateral caudate nucleus successfully discriminated familiar from missed abstract words (accuracy 75%, $p$ = 0.01) but did not significantly discriminate familiar concrete from missed concrete words (accuracy 61%, $p$ = 0.14). Consistently with the univariate effect, the significant classification in the caudate nucleus for familiar abstract words was more pronounced within the left nucleus (Left: accuracy = 80.6%, $p$ = 0.004; Right: accuracy = 63.9%, $p$ = 0.11). It is also worth noting that there was a trend for successful discrimination of familiar concrete words (versus misses) in the left caudate (accuracy = 69.44%, $p$ = 0.056), but not in the right nucleus (accuracy = 55.6%, $p$ = 0.44).

Within the parahippocampal cortex activity did not significantly discriminate familiar abstract from missed abstract words (accuracy = 63.9%, $p$ = 0.11; Fig. 5d), but it did discriminate familiar concrete from missed concrete words (accuracy = 72.2, $p$ = 0.01; Fig. 5c). The effect was more pronounced within the left (accuracy = 83.3%, $p$ = 0.001) than the right parahippocampal cortex (accuracy = 69.4%, $p$ = 0.054). It is finally worth noting that there was a trend for reliable classification accuracy for familiar versus missed abstract words within the left (but not the right) parahippocampal cortex (accuracy = 69.4%, $p$ = 0.056; Fig. 5c).

When examining the classification performance for all abstract and concrete words (irrespective of behavioural response), classification accuracy within the caudate nucleus and the parahippocampal cortex was at (or below) chance without any statistically significant classification outcome (Fig. 5d, f). The activation response within these regions, therefore, was driven predominantly by familiarity memory and not by the degree of abstractness.

### Supplementary Figures

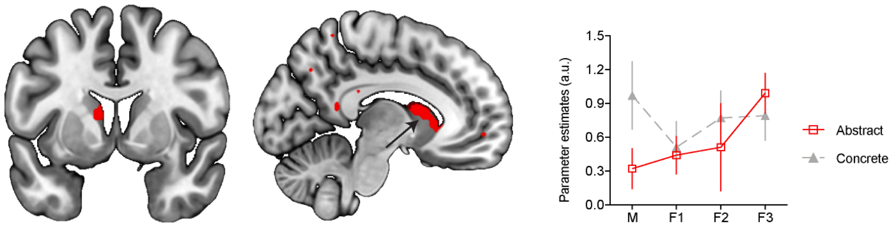


**Supplementary Fig. 1. Increases in activation as a function of familiarity memory selective for abstract words in the** caudate nucleus (MNI: -9 5 13). Activation surviving exclusive masking by concrete word familiarity is shown. Activations are displayed at a voxel-wise $p < 0.001$ and are significant at a cluster-corrected family-wise error (FWE) $p < 0.05$ determined via nonparametric permutations (all *t*s > 3.70). Note: M = misses; F1 = weak; F2 = moderate; F3 = strong familiarity. Error bars show the standard error of the mean.

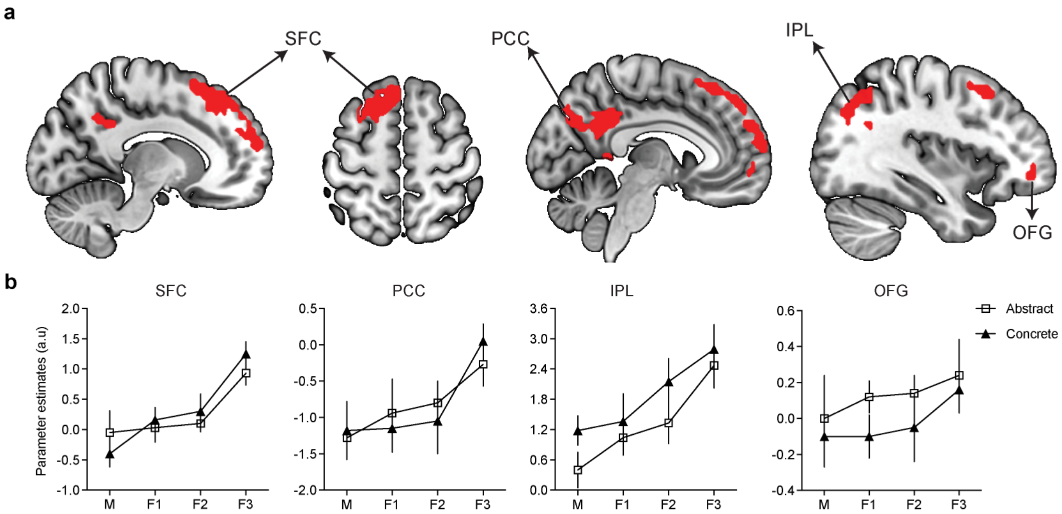


**Supplementary Fig. 2. Whole-brain responses as a function of reported familiarity memory strength** a) Whole-brain responses to abstract and concrete words across familiarity strength and b) their parameter estimates across the different memory outcomes. A conjunction analysis confirmed that these brain regions jointly responded to the degree of reported familiarity memory for both abstract and concrete words. SFC = Superior frontal cortex (BA8 and 6); OFG = orbitofrontal gyrus (BA11/47); IPL = inferior parietal lobe (BA 39 and 40); PCC = precuneus (BA7 and 21).

### Supplementary Tables

**Supplementary Table 1.** Proportion of trials, response times (RTs) and memory performance (Hits-FAs) for the different response outcomes in the recognition task separately for abstract and concrete words

| | Abstract | | | Concrete | | |
|---|---|---|---|---|---|---|
| | Proportions | Performance (H-FA) | RTs | Proportions | Performance (H-FA) | RTs |
| F1 | 0.21(0.09) | -0.10(0.13) | 1599(317) | 0.15 (0.06) | -0.08(0.13) | 1616(289) |
| F2 | 0.20(0.06) | 0.08(0.09) | 1549(227) | 0.16 (0.05) | 0.06(0.07) | 1554(269) |
| F3 | 0.38(0.13) | 0.25(0.13) | 1442(287) | 0.34 (0.15) | 0.32(0.17) | 1411(279) |
| R | 0.11(0.10) | 0.11(0.09) | 1516(312) | 0.25 (0.17) | 0.23(0.17) | 1438(228) |
| M | 0.10 (0.09) | | 1545(242) | 0.10 (0.06) | | 1507(219) |
| CR | 0.43 (0.18) | | 1552(236) | 0.63 (0.18) | | 1443(211) |
| FA F1 | 0.31 (0.09) | | 1628(311) | 0.23 (0.14) | | 1615(292) |
| FA F2 | 0.12 (0.06) | | 1562(252) | 0.10 (0.05) | | 1662(371) |
| FA F3 | 0.13 (0.11) | | 1524(419) | 0.02 (0.02) | | 1375(296) |
| FA R | 0.01 (0.01) | | 1965(462) | 0.02 (0.01) | | 1563(652) |

*Note:* Standard deviations presented in parentheses; F1= weak familiarity, F2 = moderate familiarity, F3 = strong familiarity, R = recollection, CR = correct rejections, M = misses, FA = false alarms

**Supplementary Table 2.** Parametric responses (monotonic increases) to abstractness and concreteness across the whole brain

| Side | Region | No. Voxels | BA | MNI x y z | *T*-value |
|---|---|---|---|---|---|
| *BOLD increases with increased abstractness* | | | | | |
| L | Hippocampus | 25 | | -24 -40 1 | 7.60 |
| R | Hippocampus | 9 | | 30 -25 -14 | 6.2 |
| L | Middle occipital gyrus/ Lingual gyrus | 44 | 19 | -24 -73 4 | 6.2 |
| L | Middle cingulate gyrus | 41 | 31 | -15 -31 43 | 4.42 |
| L | Insula | 50 | 13 | -39 -16 -2<br>-39 -7 1 | 4.29 |
| R | Insula | 35 | 13 | 42 -4 -5 | 4.15 |
| L | Middle cingulate | 28 | 31 | -9 -10 46 | 3.75 |
| *BOLD increases with increased concreteness* | | | | | |
| L | Precuneus & posterior cingulate | 661 | 31/7 | -3 -49 13 | 8.28 |
| L | Superior medial frontal gyrus | 614 | 6/8 | 3 29 43 | 7.92 |
| L | Parahippocampal cortex | 41 | 36 | -24 -34 -23 | 5.71 |
| L | Inferior parietal lobe/ Angular gyrus | 308 | 39/40 | -33 -70 43 | 5.7 |
| R | Caudate | 19 | | 9 11 1 | 4.04 |

*Note:* Activations are FWE-corrected at the cluster level for multiple comparisons at $p < 0.05$ determined via nonparametric permutations (*t*s > 3.75).

**Supplementary Table 3**. Activations to recollected stimuli (R > M and R > F3) for abstract words

| Side | Region | No. Voxels | BA | MNI x y z | *T*-value |
|---|---|---|---|---|---|
| *R > M* | | | | | |
| L | Angular gyrus | 20 | 39 | -42 -70 34 | 5.85 |
| L | Middle frontal gyrus | 108 | 8/9 | -27 23 55 | 4.85 |
| R | Precentral gyrus | 52 | 4 | 33 -25 55 | 4.54 |
| L | Precuneus | 54 | 31/7 | -12 -52 25 | 4.5 |
| L | Superior medial frontal gyrus | 118 | 8/6/10 | -3 65 25 | 4.37 |
| L | Caudate | 47 | | -12 8 13 | 4.35 |
| L | Thalamus (pulvinar) | 26 | | -6 -25 10 | 3.98 |
| L | Hippocampus | 11 | | -30 -13 -20 | 3.96 |
| *R > F3* | | | | | |
| L | Middle frontal gyrus | 172 | 8/6 | -27 20 46 | 5.00 |
| L | Angular gyrus | 33 | 39 | -42 -70 34 | 4.47 |
| L | Caudate | 21 | | -12 17 1 | 4.22 |
| R | Putamen | 16 | | 30 -7 -2 | 4.18 |
| R | Amygdala and hippocampus | 14 | | 21 -7 -14 | 3.97 |

*Note:* Activations are FWE-corrected at the cluster level for multiple comparisons at $p < 0.05$ determined via nonparametric permutations ($t$s > 3.75).

**Supplementary Table 4.** Activations to recollected stimuli (R > M and R > F3) for concrete words

| Side | Region | No. Voxels | BA | MNI x y z | *T*-value |
|---|---|---|---|---|---|
| *R > M* | | | | | |
| L | Posterior cingulate (and retrosplenial cortex) | 298 | 29/30 | -3 -46 4 | 8.3 |
| L | Caudate, putamen and globus pallidus | 71 | | -12 20 -2 | 5.27 |
| L | Superior frontal gyrus | 440 | 8/6 | -12 26 43 | 5.68 |
| L | Hippocampus and parahippocampal cortex | 30 | | -15 -25 -17 | 5.32 |
| L | Inferior parietal lobe (supramarginal gyrus) | 84 | 40 | -42 -52 43 | 4.1 |
| L | Orbitofrontal cortex | 274 | 10 | -18 56 7 | 4.98 |
| L | Precuneus | 20 | 7 | -3 -70 37 | 4.84 |
| *R > F3* | | | | | |
| L | Superior medial frontal gyrus and medial orbitofrontal cortex | 581 | 10/9/32 | -9 56 19 | 6.1 |
| L | Caudate, putamen | 113 | | -12 20 1 | 6 |
| L | Posterior cingulate (and retrosplenial cortex) | 136 | 30/31 | -6 -46 31 | 5.41 |
| L | Inferior parietal lobe (supramarginal gyrus) | 40 | 40 | -36 -55 25 | 4.95 |
| R | Superior temporal gyrus | 39 | 39 | 48 -58 19 | 4.04 |
| R | Caudate (head) | 13 | | 9 17 -5 | 3.94 |
| L | Hippocampus | 15 | | -24 -28 -5 | 3.95 |

*Note:* Activations are FWE-corrected at the cluster level for multiple comparisons at $p < 0.05$ determined via nonparametric permutations (*t*s > 3.75).

**Supplementary Table 5.** MVPA classification accuracy (%) at the group level for recollections (R) relative to misses (M) and F3 responses within the anatomical area of the hippocampus bilaterally (B) and separately for left (L) and right (R) masks.

| | **R vs M** | | | | **R vs F3** | | | |
|---|---|---|---|---|---|---|---|---|
| | **Abstract** | | **Concrete** | | **Abstract** | | **Concrete** | |
| | **Accuracy** | ***p*** | **Accuracy** | ***p*** | **Accuracy** | ***p*** | **Accuracy** | ***p*** |
| **B** | 67.65% | 0.056 | 70.59% | 0.04 | 67.65% | 0.054 | 73.53% | 0.03 |
| **L** | 70.59% | 0.035 | 73.53% | 0.03 | 61.76% | 0.164 | 76.47% | 0.02 |
| **R** | 67.65% | 0.054 | 70.59% | 0.05 | 70.59% | 0.05 | 67.65% | 0.054 |

*Note: p*-values indicating significance of classification success were calculated from permutation testing with 5000 permutations.

**Supplementary Table 6.** Brain activations (parametric responses) to familiar abstract and concrete words and conjunction analysis

| Side | Region | No. Voxels | BA | MNI x y z | *T*-value |
|---|---|---|---|---|---|
| *Parametric familiarity: Abstract words* | | | | | |
| L | Superior medial frontal gyrus & orbitofrontal cortex | 960 | 8/47 | -6 65 16 | 8.75 |
| L&R | Precuneus | | 7/21 | 0 -73 37 | 7.67 |
| L | Posterior cingulate cortex | 1057 | 30/31 | -3 -52 28 | 7.47 |
| L | Angular gyrus | | 39 | -39 -73 37 | 7.12 |
| L | Caudate | 55 | | -9 5 13 | 5.93 |
| L | Hippocampus | 25 | | -30 -22 -14 | 4.16 |
| *Parametric familiarity: Concrete words* | | | | | |
| L&R | Posterior cingulate | 115 | 30/31 | -3 -43 31 | 8.32 |
| L | Superior medial frontal gyrus | 743 | 8/6/10 | -9 65 22 | 6.58 |
| L | Inferior frontal gyrus & Orbitofrontal cortex | 60 | 10/11/47 | -39 47 -5 | 6.53 |
| L | Angular gyrus | 40 | 39 | -48 -58 40 | 5.22 |
| L | Inferior parietal lobe | 67 | 40 | -30 -76 43 | 5.34 |
| L&R | Precuneus | 44 | 7 | -3 -73 34 | 4.68 |
| L | Parahippocampal cortex | 22 | 35 | -22 -27 -20 | 4.45 |
| *Conjunction* | | | | | |
| L | Superior medial and lateral frontal gyrus | 596 | 8 | -12 23 58 | 6.29 |
| L | Posterior cingulate and precuneus | 234 | 7/31 | -6 -46 31 | 5.89 |
| L | Inferior parietal lobe | 142 | 40 | -36 -64 49 | 5.32 |
| L | Lateral and medial orbitofrontal cortex | 49 | 10/11 | -24 50 -2 | 4.42 |

Note: Activations are FWE-corrected at the cluster level for multiple comparisons at $p < 0.05$ determined via nonparametric permutations (*t*s > 3.70).
